# Degradation and Stable Maintenance of AAV Inverted Terminal Repeats in *E. coli*

**DOI:** 10.1101/2024.06.03.597121

**Authors:** Marco T. Radukic, Dinh To Le, Timo Krassuski, Philipp Borchert, David R. F. Leach, Kristian M. Müller

## Abstract

Current plasmid propagation compromises large inverted repeats, like inverted terminal repeats (ITRs) of adeno-associated virus (AAV). Direct long-read sequencing analyses upon varying culture conditions and strains revealed ITR instability mechanisms, which diminished in absence of SbcC or at elevated growth temperatures (e.g. 42 °C), with a combination being optimal. Resulting full ITRs improved rAAV yield and purity. The findings advance plasmid biology, cloneable sequences, and therapeutic AAV manufacturing.

## Main

AAV vectors have facilitated a paradigm shift in the treatment of human genetic diseases but patient safety and access demand improvements ^1,2^. The instability of AAV ITRs on plasmids during propagation in *E. coli* ^3^ poses a critical manufacturing challenge, particularly at the industry scale, where transient transfection involves gram amounts of plasmid for reactors of more than a thousand liters. ITRs are nested palindromes of about 145 bp and particularly elusive due to their sequence and multiple biological functionalities (**Figure 1a** for ITR drawing; for a review see Berns, 2020 ^4^). Conventionally, the field has opted for plasmids encoding preemptively truncated ITRs (130 bp) instead of wild-type ITRs, hoping to sidestep the instability issue ^5^. A meta-analysis of published plasmids (SI for method) showed that, of 123 sequence-verified ITR-plasmids published by the plasmid repository Addgene, none were full-length wildtype ITRs, as expected, but 69 plasmids had acquired further unintended internal deletions (119 bp – 108 bp ITRs) possibly throughout their cloning and propagation history. These findings, among others, highlight that the ITR instability challenge remains unsolved and affects pharma industry after 40 years of AAV research ^6^.

**Figure 1:**
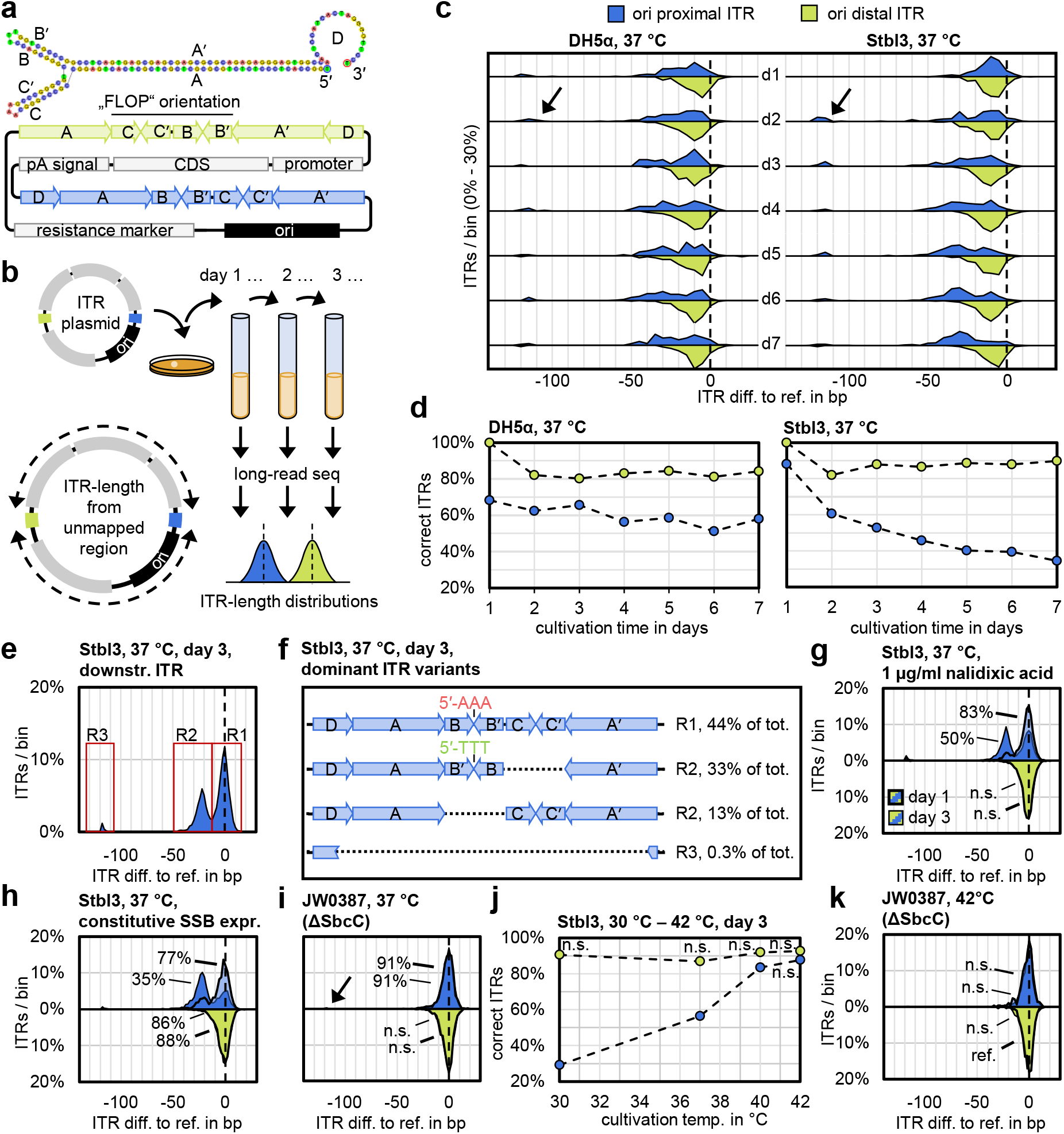
Analysis of inverted repeat stability in E. coli. **a**, ITR and plasmid scheme. **b**, single ITR sequencing method overview. **c**, ITR-length distribution histograms for a wildtype ITR plasmid (pZMB938) propagated for seven days in two different strains. **d**, Percentage of remaining correct ITRs of (c). **e**, ITR-length distribution histograms for another wildtype ITR plasmid (pZMB990) with regions of interest (R1 – 3). **f**, underlying sequences per region of (e). **g**, ITR degradation in presence of DNA gyrase inhibitor nalidixic acid. **h**, ITR degradation in presence of constitutive E. coli SSB expression. **i**, wildtype ITR stability in ΔSbcC strain JW0387. **j**, cultivation temperature-dependence of ITR stability. **k**, ITR stability in JW0387 cultivated at 42 °C. n.s.: not significant. *p < 0.05. Percentages in charts are percent correct ITRs.

We have previously characterized an internal 11 bp ITR deletion and presented evidence that the deletion contributes to genomic heterogeneity of rAAV, despite the ITR tendency to self-correct during rAAV production in mammalian cells, which motivated us to look into large inverted repeat instability in *E. coli* ^7–10^.

We relate this phenomenon to the known instability of other large inverted repeats ^11^. Such instability has been attributed to replication fork stalling at secondary structures, which can form during strand separation within the replication fork, where they compete with single-strand binding protein (SSB) ^12,13^. Inverted repeats may also be extruded by negative supercoiling to cruciform-like structures ^14–18^. Highly negatively supercoiled DNA has been suggested to originate from DNA gyrase overcompensating for the positive supercoiling in front of the fork ^19^. A detailed sequence-level understanding of the degradation products is required to further elucidate the degradation mechanism and solve the challenge.

In design and test cycles of cloning and culturing, we investigated ITR degradation by long-read sequencing of individual ITR plasmid molecules (**Figure 1a** for plasmid scheme) propagated in typical strains first under standard conditions (**Figure 1b** for method scheme, **SI Figure 1** for method qualification). Pairs of wild-type ITRs degraded with the known predominance in the ITR located in proximity to the origin of plasmid replication over three days or more, a typical timespan of a large scale plasmid production (**Figure 1c,d**). These results were consistent between plasmids harboring full-length ITRs obtained from rAAV or cloned from hybridized oligo nucleotides (see **SI Figure 2** and **SI Figure 3** for respective cloning strategies). Detailed, nucleotide-level analysis of the ITR degradation sequences of a representative dataset (**Figure 1e**) showed that ITR degradation resulted in two main outcomes: (i) internal deletion of the C hairpin and concomitant inversion of the B hairpin (59% of sequences with deletion); or (ii) internal deletion of the B hairpin while the C hairpin was not inverted (24% of sequences with deletion) (**Figure 1f** and **SI Figure 4**). Some plasmids also completely lost their inverted repeat with only a partial D- and A-sequence remaining (**Figure 1c**, black arrows, and 0.6% of sequences with deletion **Figure 1e**). ITR degradation prevailed also when we constitutively overexpressed the *E. coli* SSB or inhibited DNA gyrase by viable concentrations of nalidixic acid (**Figure 1g,h**).

In a common model of inverted repeat instability on plasmids, *E. coli* SbcCD, a homolog of the human MRN complex Mre11/RAD50, incises DNA hairpins and possesses 3’→5’ exonuclease activity ^20^. This action exerts selective pressure against inverted repeats by degrading potential secondary structures, which leads to subsequent plasmid loss by the RecBCD exonuclease. The pressure is relieved by acquiring mutations via a “slipped misalignment” direct repeat contraction ^21^. Indeed, a ΔSbcC strain, JW0387, showed a vast improvement in ITR stability (**Figure 1i**). However, complete loss of the downstream ITR remained in some cases (**Figure 1i**, black arrow), in line with reports of the instability of large, albeit TA-rich, inverted repeats in the low-yield ΔSbcC strain SURE2 ^22–24^. ITR degradation also occurred when only one ITR was present per plasmid (**SI Figure 2**) though cloning of fragments with two ITRs under standard conditions was particularly ineffective.

Since degradation appeared to involve secondary-structure formation and enzymatic cleavage we tested the influence of cultivation temperature on ITR stability. ITRs stabilized with increased temperature (**Figure 1j**) both in terms of internal degradation and complete ITR loss. Stabilization was observed for all tested strains to varying degrees (**SI Figure 5**) and was most prominent in the ΔSbcC strain, where ITR degradation at 42 °C was insignificant (**Figure 1k**). The increased temperature also did not influence plasmid quality at useful yields and facilitated full ITR cloning even in standard cloning strains (**SI Figure 6** and **SI Table 1**). These findings essentially solve the ITR instability challenge. We assume that the increased temperature helps to avoid and resolve undesired DNA hybridization but also indirect effects on temperature sensitive proteins and/or upregulation of host factors may play a role.

We could rationalize the specific ITR degradation products (**Figure 1f**) with a slippage model. Both, the simple deletion and the deletion-inversion appear to rely on a slippage at a CG dinucleotide direct repeat, while the deletion-inversion additionally requires back-amplification of the counter-strand (**SI Figure 4** for sequences and **SI Figure 7** for model). The complete ITR loss appears to occur at a GA direct repeat. Still, slippage cannot account for the elevated instability of the ori proximal ITR or the dominance of the observed deletions.

Additional mechanisms may contribute. For example, the ITR D-A′ interface 5′-GTTG-3′ bears similarity to the Holliday-resolvase RuvC consensus cleavage site 5’-(A/T)TT(C/G)-3’ ^25^. We speculate that an initial high level of positive supercoiling near the origin of replication, when the replication bubble is first established, and overcorrection by DNA gyrase, could extrude ITRs near the ori to Holliday-like structures and predominantly destabilize this ITR. Isomerization of these structures would lead to non-replicative hairpin inversion (**SI Figure 8** for model). Resulting closed-ended intermediates then share terminal homologies amenable to homologous RecA-independent repair, with the specific observed deletion depending on the extent of exonuclease digestion prior to joining ^26,27^. The near-complete ITR loss would be facilitated, in this sense, by a partial homology between the D and A sequence, which forms the ITR terminal resolution site ^28^.

Interestingly, since ITR deletions have also been observed in AAV genome concatemers formed in mammalian cells, further elucidation of the ITR degradation mechanisms across hosts is expected to inform improvements to sustained AAV-based transgene expression in gene therapy ^29,30^.

Finally, our findings enabled us to prepare plasmids containing truly stable, full-length ITRs, to test in rAAV production. When we compared rAAV production from truncated but otherwise non-deleted ITRs (130 bp length), to rAAV production with one truncated and one additionally internally deleted ITR (119 bp length), a slight increase in titer from the solely truncated ITR was accompanied by a large and significant increase in genomic purity (**Figure 2a,b**). When using plasmids containing stably maintained full-length ITRs (143 bp, 145 bp), titers and genomic purity again increased significantly. In addition, the genome-to-capsid ratio also appeared to increase with ITR integrity (**Figure 2c**). These findings support our initial assessment that the genome correction mechanism during production takes time and is error-prone, giving room for mispackaging events. Upon transfection of the wildtype ITR bearing plasmids, rAAV genome rescue and replication appear facilitated, leading to more and purer product, a measure that should be easily adopted in any common AAV production process, even at large scale without further changes to bulk materials, equipment, and operating procedures.

**Figure 2:**
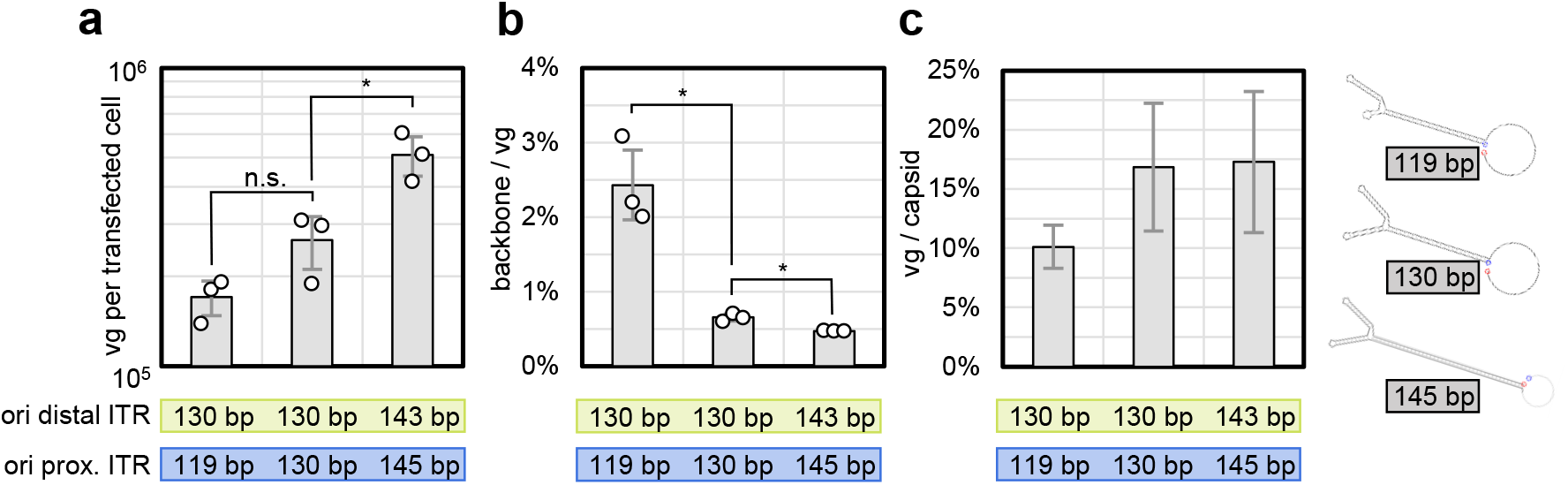
rAAV production from different ITR variants. **a**, viral genomes (vg) per cell. **b**, mispackaged DNA as fraction backbone to vg. **c**, fraction “full” capsids. n.s.: not significant. *p < 0.05.

In conclusion, we show that the complex issue of inverted repeat instability, exemplified with inverted terminal repeats of AAV, can be solved by the simple measure of propagating *E. coli* ΔSbcC at 42 °C. Importantly, we demonstrate that an increased growth temperature is a key factor for the maintenance of AAV2 ITRs, without compromising sequence quality. This worked across various *E. coli* strains for cloning and plasmid production. The immediate practical implication is an improvement in rAAV production through the use of full-length ITRs, one vital building block in fulfilling the promises of genetic medicine for all patients. In a broader sense, our findings should enable cloning and maintenance of DNA constructs typically foreign to plasmids but useful in synthetic biology applications, like DNA frameworks for nanotechnology and synthetic viral genomes.

## Methods

### E. coli culture and strains

Ca^2+^-chemically competent *E. coli* were transformed and plated at the temperature indicated in the main text and incubated on selection plates overnight. Colonies were picked into 5 ml lysogeny broth (LB, Lennox) in test tubes and cultured overnight at the indicated temperature. For continued cultivation, each morning, 10 μl overnight culture were transferred to 5 ml fresh broth and the culture was again diluted in the evening. *E. coli* DH5alpha was from DSMZ (No. 6897). Stbl3 was from Thermo. BL21(DE3) was from DSMZ (No. 102052). JW0387 was from the Keio collection (National Institute of Genetics, Japan) ^31^. JM103 was from DSMZ (No. 2829). Ampicillin concentration was 100 μg/ml (37 °C) and 300 μg/ml (42 °C).

### Molecular cloning

pZMB938 is a pUC19-based plasmid carrying an mVenus fluorescence reporter transgene and ori distal a 143 bp ITR and ori proximal a 145 bp ITR, which were recovered from an rAAV source (**SI Figure 9** for plasmid map). rAAV were produced, purified by affinity chromatography and the DNA was extracted as previously described ^8^. Hybridized genomes were cut with SacI and separated on an agarose gel. The two transgene fragments representing the upstream and downstream segments of the recombinant genome were separately ligated with a pUC19 backbone opened with SacI and HincII. Colonies were then screened for full ITRs using Nanopore sequencing consensus analysis. A transgene upstream segment containing a 143 bp ITR was excised with SacI and PvuII and cloned into a 145 bp ITR containing plasmid (downstream segment) opened with SacI and SfoI, thereby reconstituting the complete transgene.

Plasmid pZMB990 is a pUC19-based plasmid carrying an eGFPd2 fluorescence reporter flanked by two 144 bp ITRs, which were obtained from hybridized oligonucleotides (**SI Figure 10** for plasmid map). A scaffolding plasmid to accept the ITR-oligos was first constructed. For this, the PciI recognition site of a pUC19 plasmid was knocked out by PciI digest, blunting and religation. The multicloning site of the plasmid was then excised with EcoRI and PstI and a new multi-cloning site containing SalI, PstI, MlyI, NcoI, EcoRI, SpeI, PciI, MlyI(inv), PstI, and XhoI in this order was inserted from synthetic oligos. SalI and XhoI, as well as NcoI and PciI, produce compatible cohesive ends. Six phosphorylated oligonucleotides comprising the full-length 144 bp AAV serotype 2 ITR sequence with overhangs for ligation to SalI and NcoI overhangs and already containing the MlyI and PstI sites of the novel multi-cloning sites were then hybridized and ligated. The ligation product was separated on an agarose gel (1%, TAE, 120 V, 50 min.), the band of the expected size was cut out, and the DNA was purified. The scaffolding plasmid was cut with SalI and NcoI, and the backbone was ligated with the purified ITR fragment. *E. coli* Stbl3 were transformed with the ligation product and plated and cultured at 42 °C. A transgene encoding eGFPd2 was then cloned into the intermediate plasmid using the EcoRI and SpeI restriction sites. The downstream ITR was then excised with SalI and NcoI and purified by gel extraction. Separately, the intermediate plasmid was digested with PciI and XhoI. The ITR fragment was then ligated with the prepared backbone, resulting in plasmid pZMB990. All propagation and cloning procedures based on these plasmids were done at 42 °C.

Where applicable, strains were adapted to growth at 42 °C prior to preparation of chemically competent stocks.

### Direct single-molecule ITR sequencing

Plasmid DNA for sequencing was extracted using a plasmid kit (Macherey-Nagel NucleoSpin Plasmid mini) as per the manufacturer’s instructions. Initially, the Oxford Nanopore SQK-RBK004 sequencing kit was used with sequencing on an R9.4.1 Flongle. For this approach, 200 ng DNA per sample in 4.5 μl water in a PCR tube were incubated between thumb and index finger with 0.5 μl barcoding mix under constant pipette stirring for exactly 30 s and immediately heat-inactivated at 80 °C for exactly 1 min. This protocol adaptation was found to produce mostly full-length plasmid reads. Further library preparation, Flongle loading, and sequencing was as per manufacturer’s instructions. Later in the study, the updated Oxford Nanopore kit SQK-RBK114 with sequencing on R10.4.1 Flongle flow cells was used. For this approach, 50 ng DNA per sample in 4.5 μl water in a PCR tube were incubated between thumb and index finger with 0.5 μl barcoding mix under constant pipette stirring for 30 s and immediately heat-inactivated at 80 C for 2 min, otherwise following the manufacturer’s instructions.

### Bioinformatic analysis

R9 reads were basecalled using ONT Guppy (6.5.7). R10 reads were basecalled using ONT Dorado (0.4.3). Consensus sequences were obtained using Flye (2.8.3) for draft assembly and Medaka (1.6.0 consensus wrapper) for polishing. Base methylations were called using Dorado with the 4.2.0 models (5mC and 6mA). ITR length per single read were determined using a custom Python script (see supplementary information) built around mappy (2.24) and pandas (2.0.3). ITR-lengths populations were compared using a t-test (two-sided, heteroscedastic test). The percent-degradation between significantly different (p < 0.05) populations was calculated by binning the distributions (2 bp or 5 bp bin width for R10 and R9 reads, respectively), normalizing the total number of analyzed ITRs per distribution to one and subtracting the respective bins of the normalized histograms.

### rAAV production analysis

rAAV were produced as we have previously described ^8^, except that all productions were downscaled by growth area to 6-well plate format with each well constituting a biological replicate treated separately. Briefly, adherent HEK-293 cells were triple-transfected (pHelper, pRepCap, and pITR-GOI) and rAAV were harvested after three days of incubation. rAAV were purified from the lysate by AAVX affinity purification (Thermo) as batch purification, and the genomic titer (CMV promoter) and backbone titer (*beta lactamase* gene) were determined by qPCR with the previously qualified primer sets.

## Acknowledgements

We thank Dr. Julian Teschner for early contributions to cloning procedures and Daniel Falk for contributing to experiments. We thank the NBRP-*E*.*coli* at the National Institute of Genetics, Japan, for providing strain JW0387. MTR acknowledges support by the Bielefeld University Young Researcher’s Fund. DTL acknowledges Deutscher Akademisher Austauschdienst (DAAD) for providing a fellowship (personal ref no. 91651068).

## References

1. Sheridan, C. Why gene therapies must go virus-free. Nat. Biotechnol. 41, 737–739 (2023).

2. Shen, W., Liu, S. & Ou, L. rAAV immunogenicity, toxicity, and durability in 255 clinical trials: A meta-analysis. Front. Immunol. 13, 1001263 (2022).

3. Samulski, R. J., Berns, K. I., Tan, M. & Muzyczka, N. Cloning of adeno-associated virus into pBR322: rescue of intact virus from the recombinant plasmid in human cells. Proc. Natl. Acad. Sci. 79, 2077–2081 (1982).

4. Berns, K. I. The Unusual Properties of the AAV Inverted Terminal Repeat. Hum. Gene Ther. 31, 518–523 (2020).

5. Samulski, R. J., Chang, L. S. & Shenk, T. A recombinant plasmid from which an infectious adeno-associated virus genome can be excised in vitro and its use to study viral replication. J. Virol. 61, 3096–3101 (1987).

6. Chen, Y. et al. A Comprehensive Study of the Effects by Sequence Truncation within Inverted Terminal Repeats (ITRs) on the Productivity, Genome Packaging, and Potency of AAV Vectors. Microorganisms 12, 310 (2024).

7. Feiner, R. C. et al. rAAV Engineering for Capsid-Protein Enzyme Insertions and Mosaicism Reveals Resilience to Mutational, Structural and Thermal Perturbations. Int. J. Mol. Sci. 20, 5702 (2019).

8. Radukic, M. T., Brandt, D., Haak, M., Müller, K. M. & Kalinowski, J. Nanopore sequencing of native adeno-associated virus (AAV) single-stranded DNA using a transposase-based rapid protocol. NAR Genomics Bioinforma. 2, lqaa074 (2020).

9. Samulski, R. J., Srivastava, A., Berns, K. I. & Muzyczka, N. Rescue of adeno-associated virus from recombinant plasmids: Gene correction within the terminal repeats of AAV. Cell 33, 135–143 (1983).

10. Musatov, S., Roberts, J., Pfaff, D. & Kaplitt, M. A cis -Acting Element That Directs Circular Adeno-Associated Virus Replication and Packaging. J. Virol. 76, 12792–12802 (2002).

11. Lilley, D. M. J. In vivo consequences of plasmid topology. Nature 292, 380–382 (1981).

12. Azeroglu, B., Lincker, F., White, M. A., Jain, D. & Leach, D. R. F. A perfect palindrome in the Escherichia coli chromosome forms DNA hairpins on both leading-and lagging-strands. Nucleic Acids Res. 42, 13206–13213 (2014).

13. Voineagu, I., Narayanan, V., Lobachev, K. S. & Mirkin, S. M. Replication stalling at unstable inverted repeats: Interplay between DNA hairpins and fork stabilizing proteins. Proc. Natl. Acad. Sci. 105, 9936–9941 (2008).

14. Eykelenboom, J. K., Blackwood, J. K., Okely, E. & Leach, D. R. F. SbcCD Causes a Double-Strand Break at a DNA Palindrome in the Escherichia coli Chromosome. Mol. Cell 29, 644–651 (2008).

15. Benham, C. J. Stable cruciform formation at inverted repeat sequences in supercoiled DNA. Biopolymers 21, 679–696 (1982).

16. Matek, C., Ouldridge, T. E., Levy, A., Doye, J. P. K. & Louis, A. A. DNA Cruciform Arms Nucleate through a Correlated but Asynchronous Cooperative Mechanism. J. Phys. Chem. B 116, 11616–11625 (2012).

17. Mizuuchi, K., Mizuuchi, M. & Gellert, M. Cruciform structures in palindromic DNA are favored by DNA supercoiling. J. Mol. Biol. 156, 229–243 (1982).

18. Shlyakhtenko, L. S., Potaman, V. N., Sinden, R. R. & Lyubchenko, Y. L. Structure and dynamics of supercoil-stabilized DNA cruciforms. J. Mol. Biol. 280, 61–72 (1998).

19. Lai, P. J. et al. Long inverted repeat transiently stalls DNA replication by forming hairpin structures on both leading and lagging strands. Genes Cells 21, 136–145 (2016).

20. Gut, F. et al. Structural mechanism of endonucleolytic processing of blocked DNA ends and hairpins by Mre11-Rad50. Mol. Cell 82, 3513-3522.e6 (2022).

21. Connelly, J. C. & Leach, D. R. F. The sbcC and sbcD genes of Escherichia coli encode a nuclease involved in palindrome inviability and genetic recombination. Genes Cells 1, 285–291 (1996).

22. Palmer, E. L., Gewiess, A., Harp, J. M., York, M. H. & Bunick, G. J. Large-Scale Production of Palindrome DNA Fragments. Anal. Biochem. 231, 109–114 (1995).

23. Inagaki, H. et al. Palindromic AT-rich repeat in the NF1 gene is hypervariable in humans and evolutionarily conserved in primates. Hum. Mutat. 26, 332–342 (2005).

24. Kogo, H. et al. Cruciform extrusion propensity of human translocation-mediating palindromic AT-rich repeats. Nucleic Acids Res. 35, 1198–1208 (2007).

25. Górecka, K. M. et al. RuvC uses dynamic probing of the Holliday junction to achieve sequence specificity and efficient resolution. Nat. Commun. 10, 4102 (2019).

26. Coniey, E. C., Saunders, V. A., Jackson, V. & Saunders, J. R. Mechanism of Intramolecular recyclizatlon and deletion formation following transformation of Escherichia coli with linearized plasmid DNA. Nucleic Acids Res. 14, 8919–8932 (1986).

27. Nozaki, S. & Niki, H. Exonuclease III (XthA) Enforces In Vivo DNA Cloning of Escherichia coli To Create Cohesive Ends. J. Bacteriol. 201, (2019).

28. Brister, J. R. & Muzyczka, N. Mechanism of Rep-mediated adeno-associated virus origin nicking. J. Virol. 74, 7762–7771 (2000).

29. Duan, D., Yan, Z., Yue, Y. & Engelhardt, J. F. Structural Analysis of Adeno-Associated Virus Transduction Circular Intermediates. Virology 261, 8–14 (1999).

30. Nowrouzi, A. et al. Integration frequency and intermolecular recombination of rAAV vectors in non-human primate skeletal muscle and liver. Mol. Ther. 20, 1177–1186 (2012).

31. Baba, T. et al. Construction of Escherichia coli K-12 in-frame, single-gene knockout mutants: the Keio collection. Mol. Syst. Biol. 2, 2006.0008 (2006).

