## Supplementary Information for "Degradation and Stable Maintenance of AAV Inverted Terminal Repeats in *E. coli*"

### 1 Meta analysis of ITR degradation status in Addgene sequence-verified plasmids

The sequences of all 188 ITR-encoding plasmids (at the time of analysis) were programmatically downloaded from the Addgene database (Addgene.org, accessed 17 January 2023, plasmids satisfying the search terms "AAV" and "ITR"). All 123 sequences marked by Addgene as "sequence verified" were saved for further analysis (Addgene IDs 104839, 105669, 105677, 105678, 105679, 113763, 120219, 121502, 121921, 125713, 126520, 126521, 129267, 135964, 135965, 135966, 135967, 137188, 139979, 139980, 154849, 154871, 160069, 163791, 163792, 163793, 163794, 163795, 163796, 163797, 164468, 165441, 166870, 166871, 166872, 166873, 167577, 167578, 167579, 167580, 167581, 170117, 170118, 170120, 170121, 170122, 170123, 174703, 180183, 180184, 180191, 180192, 180193, 183078, 183079, 183081, 184002, 184003, 186418, 186419, 187181, 187182, 190473, 190474, 190475, 191523, 191795, 192569, 192570, 192572, 192573, 192575, 192576, 193166, 193168, 193169, 193170, 193171, 193172, 193174, 193175, 193176, 193177, 193178, 193179, 193180, 193181, 193182, 193183, 193185, 193186, 193187, 193188, 26968, 26969, 26971, 26972, 26973, 26975, 26976, 50455, 50464, 50466, 60226, 60227, 60229, 65417, 65418, 82705, 85451, 86949, 86950, 86954, 86955, 87951, 89256, 89570, 98216, 98217, 98218, 98219, 98220, 98704) using the following Python code:

```
import requests as rq
import time
from bs4 import BeautifulSoup

base_url = 'https://www.addgene.org/'
addgene_seqs = {}

# A function to download Addgene sequences
def get_seq(url):
    response = rq.get(url)
    response.raise_for_status()
    soup = BeautifulSoup(response.content, 'html.parser')
    return soup.find('textarea').get_text()

# A text file containing Addgene ID's to download
with open("ITR-plasmids.txt", "r") as f:
    addgene_ids = f.read().split('\n')

# Downloading sequences
for addgene_id in addgene_ids:
    url = base_url + addgene_id + '/sequences/'
    addgene_seqs[addgene_id]=get_seq(url)
    time.sleep(.1)

# Determine if the plasmid has been NGS verified
def is_NGS_verified(__string):
    if __string == '> Addgene NGS Result':
        return True
    else:
        return False

# Parse sequence
for addgene_id in addgene_seqs:
    addgene_seqs[addgene_id] = (is_NGS_verified(addgene_seqs[addgene_id][:20]),
addgene_seqs[addgene_id])

for addgene_id in addgene_seqs:
    try:
        addgene_seqs[addgene_id] = (addgene_seqs[addgene_id][0],
addgene_seqs[addgene_id][1].split('\n', 1)[1].replace('\n', ''))
    except:
        print('nothing parsed')

# Save results
with open("addgene_itr_plasmids.fasta", "w") as fasta_file:
    for addgene_id in addgene_seqs:
        if addgene_seqs[addgene_id][0] == True:
            fasta_file.write(">" + addgene_id + '_verified' + "\n" +
addgene_seqs[addgene_id][1] + "\n")
        else:
            fasta_file.write(">" + addgene_id + '_unverified' + "\n" +
addgene_seqs[addgene_id][1] + "\n")
```

Downloaded sequences were then mapped to the AAV2 ITR reference sequences (in FLIP and FLOP configuration:

```
import mappy as mp
import pandas as pd

# A fasta file containing wildtype AAV2 ITR sequences in FLIP and FLOP config.
# A fasta file containing Addgene ITR plasmids
ref="ITR-refs.fasta"
sequin="addgene_itr_plasmids.fasta"

# mapping is done using minimap2 mappy
a = mp.Aligner(ref, scoring=[2, #Matching Score [2] [default map-ont]
                             4, #Mismatching penalty [4]
                             4, #Gap open penalty [4]
                             2, #Gap extension penalty [2]
                             24,#Long gap open penalty [24]
                             1, #Long gap extension penalty [1]
                             ]) #load or build index
if not a: raise Exception("ERROR: failed to load/build index")

data = []
columns = ["seq_id",      #unique read identifier (given by sequencer)
            "verified",
            "q_en",       #end of alignment on querey (read)
            "seq_len",    #length of read
            "map_len",    #length of mapping
            "matches",    #number matching bases
            "ctg",        #name of reference used for alignment
            "r_st",       #start of alignment on reference
            "r_en",       #end of alignment on reference
            "q_st",       #start of alignment on query
            "strand"]

for name, seq, qual in mp.fastx_read(sequin): # read a fasta/q sequence
    for hit in a.map(seq):
        tmp = [name.split('_')[0], name.split('_')[1], hit.q_en,  len(seq),
hit.blen, hit.mlen, hit.ctg, hit.r_st, hit.r_en, hit.q_st, hit.strand]
        data.append(tmp)
df = pd.DataFrame(data, columns=columns)
df = df.sort_values(by=["seq_id", "q_st"], ignore_index=True)
df_grouped = df.groupby(["seq_id", "q_st"], sort=False).last()
df_grouped
```

Further analysis could then be performed by querying of the resulting DataFrame. To determine the number of sequence-verified ITRs (with correct D-sequence to exclude plasmids for scAAV production) we used:

```
len(df_grouped.loc[(df_grouped['verified'] == 'verified') &
(df_grouped['r_en'] == 145)].index.unique(level=0))
```

The number of plasmids containing ITRs longer than 130 bp was determined with:

```
len(df_grouped.loc[(df_grouped['verified'] == 'verified') &
(df_grouped['matches'] > 130)].index.unique(level=0))
```

The number of plasmids containing ITRs with internal deletions was determined with:

```
len(df_grouped.loc[(df_grouped['verified'] == 'verified') &
(df_grouped['matches'] < df_grouped['map_len'])].index.unique(level=0))
```

### 2 ITR-length measurement method qualification

We developed a Nanopore sequencing workflow to quantify ITR degradation. Nanopore technology is capable of direct sequencing with each sequencing read representing one plasmid molecule within a population. Our workflow determines the length of an ITR as the number of basecalled bases between the ITR-adjacent sequences (e.g. the transgene and the plasmid backbone). The called lengths will follow a normal distribution even for stable sequences because of the given precision of the sequencing technology. These distributions can then be compared by common statistical tools to determine the extent of ITR degradation. We qualified the workflow for the available nanopore R9.4.1 (abbreviated R9, **SI Figure 1a – c**) and the most recent pore R10.4.1 (abbr. R10, **SI Figure 1d – f**) by sequencing an available ITR plasmid (lab. ref. pZMB522) carrying transgene upstream a 130 bp ITR (**SI Figure 1**, green color) and downstream, proximal to the ori, a 119 bp ITR (**SI Figure 1**, blue color) with an 11 bp internal deletion. We then calculated the ITR length for each sequenced plasmid molecule separately and compared the ITR length distributions between the two ITRs and the two nanopore chemistries.

We found that the R10 pore gives superior accuracy for a 130 bp ITR sequence compared to the R9 pore, while the ITR length-call precision is also generally improved, evident by the lower peak width and peak mean localization of the R10 dataset. For further method qualification, artificially mixed read populations of the 119 bp and 130 bp ITR were produced and compared to the original 130 bp ITR population using a t-test and by calculating an ITR-degradation percentage (normalized histogram subtraction). The higher precision and accuracy of the R10 pore resulted in a lower limit of degradation detection. Comparing the expected degradation percentage with the calculated degradation allows to determine the systematic error of the method, which was 34% for the R9 pore and 19.1% for the R10 pore. Since the systematic error is dependent on the absolute base loss, which could be more than the qualified 11 bp, we opted not to correct for it throughout this work. We used both pore chemistries as available to us at the time of analysis as we could show that both pores are principally suitable to resolve ITR degradation.

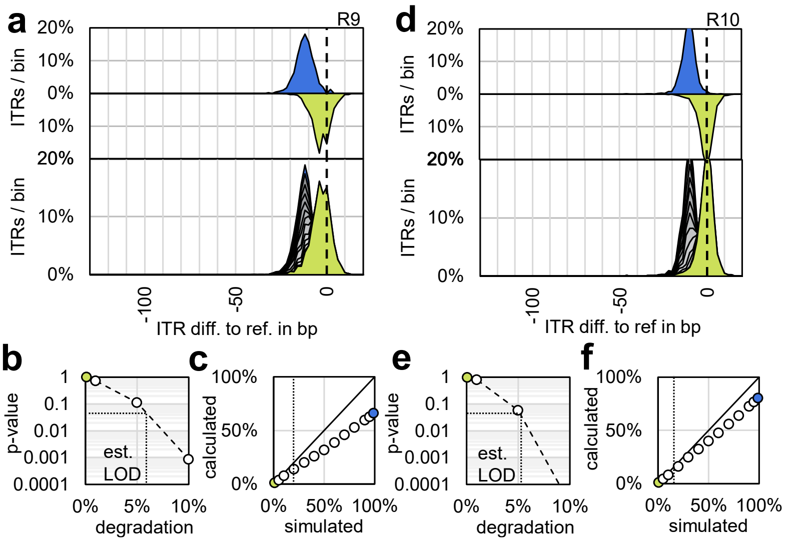

***SI Figure 1:*** *Qualification of the Nanopore method for ITR-length degradation analysis.* ***a****, plasmid containing upstream a 130 bp ITR and downstream a 119 bp ITR (lab ref. pZMB522) was sequenced with R9 kit chemistry (****a*** *-* ***c****) and R10 kit chemistry (****d*** *-* ***f****). Histograms of ITR length distributions as difference to the expected ITR length (130 bp) are shown (****a****,* ***d****, upper two distributions). The underlying length were computationally mixed at different ratios to simulate ITR degradation (****a****,* ***d****, lower distribution). The limit of detection (panels* ***b****,* ***e****) and systematic error (panels* ***c****,* ***f****) for ITR degradation in this context was then estimation from these artificial mixtures. Both sequencing chemistries are principally capable to resolve ITR degradation. The R10 chemistry offers a lower limit of detection and lower systematic error compared to R9 kit chemistry.*

### 3 Comparison of E. coli strains for ITR integrity.

***Table 1:*** *Comparison of ITR stability and plasmid yield in different strains based on pZMB938 or pZMB990.*

| Strain | potentially relevant genotype | downstream ITR stab. at day 3, 37 °C | downstream ITR stab. at day 3, 42 °C | Final $OD_{600}$ | Yield in µg / 5 ml culture | dam / dcm methylation in % |
|---|---|---|---|---|---|---|
| DH5α | gyrA96, recA1, endA1, hsdR17 | 66% | 80% | n.d. | n.d. | n.d. |
| Stbl3 | recA13 | 68% | 94% | 5 ± 3 (37 °C)<br>2.5 ± 0.6 (42 °C) | 5 ± 3 (37 °C)<br>10 ± 4 (42 °C) | 72 ± 9 (dam, 37 °C)<br>64 ± 7 (dcm, 37 °C) |
| JM103 | endA1, sbcB15 | 67% | n.d. | n.d. | n.d. | n.d. |
| BL21(DE3) | dcm, hsdSB | 30% | 64% | n.d. | n.d. | n.d. |
| JW0387 | sbcC, hsdR514 | 91% | n.s. | 4 ± 2 (37 °C)<br>4 ± 2 (42 °C) | 8 ± 1 (37 °C)<br>2 ± 1 (42 °C) | 62 ± 13 (dam, 42 °C)<br>42 ± 13 (dcm, 42 °C) |

*n.s.: no significant degradation (p > 0.05). n.d.: not determined. The standard deviation is given.*

### 4 Recovery of full ITRs from rAAV produced with plasmids encoding truncated ITRs

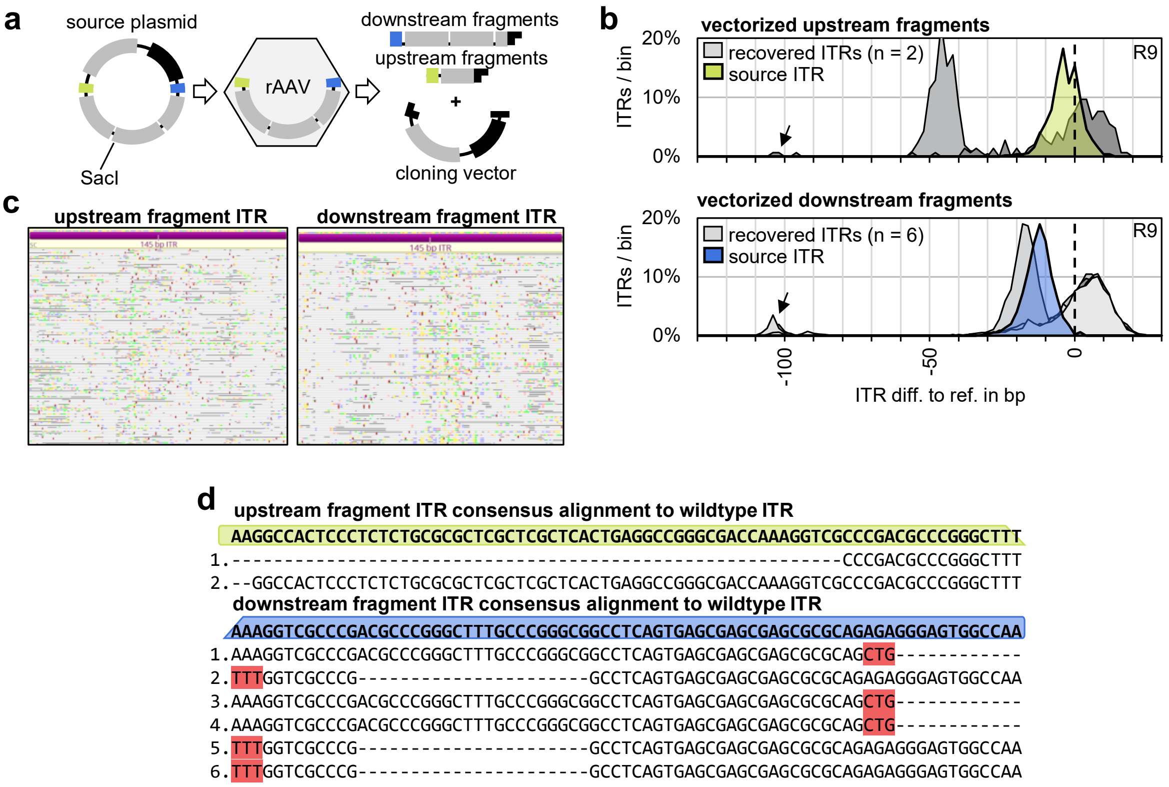

***SI Figure 2:*** *ITR cloning from rAAV. rAAV were produced from plasmids containing truncated source ITRs and full ITRs were recovered from AAV DNA as described in the main method section.* ***a****, overview of the cloning procedure as described in the main methods.* ***b****, ITR length distributions for recovered ITRs (grey) compared to the source ITRs (green: upstream ITR or ori distal ITR on the source plasmid, blue: downstream ITR or ori proximal ITR on the source plasmid) as difference to the expected size (130 bp). Results are based on R9 sequencing chemistry. Initial ITR degradation is visible for all recovered ITRs (tailing of main peak and black arrows) even though only one recovered ITR is present per plasmid in this analysis. A gain in length for some recovered ITRs compared to the respective source ITRs indicates that an ITR repair has occurred.* ***c****, multiple-read alignment from samples containing ITRs with a net gain in ITR length to the wildtype ITR reference sequence (FLOP) shows that the full ITR sequence is principally present in these samples.* ***d****, alignment of the consensus sequences of the recovered ITRs (Flye + medaka_consensus) shows mismatches, likely arising from the partial ITR deletions already present in these plasmid preparations. These findings highlight that one must be very careful when assessing ITR sequences of (partially) deleted ITRs from the consensus alone.*

### 5 Cloning of synthetic ITRs from oligonucleotides and their degradation

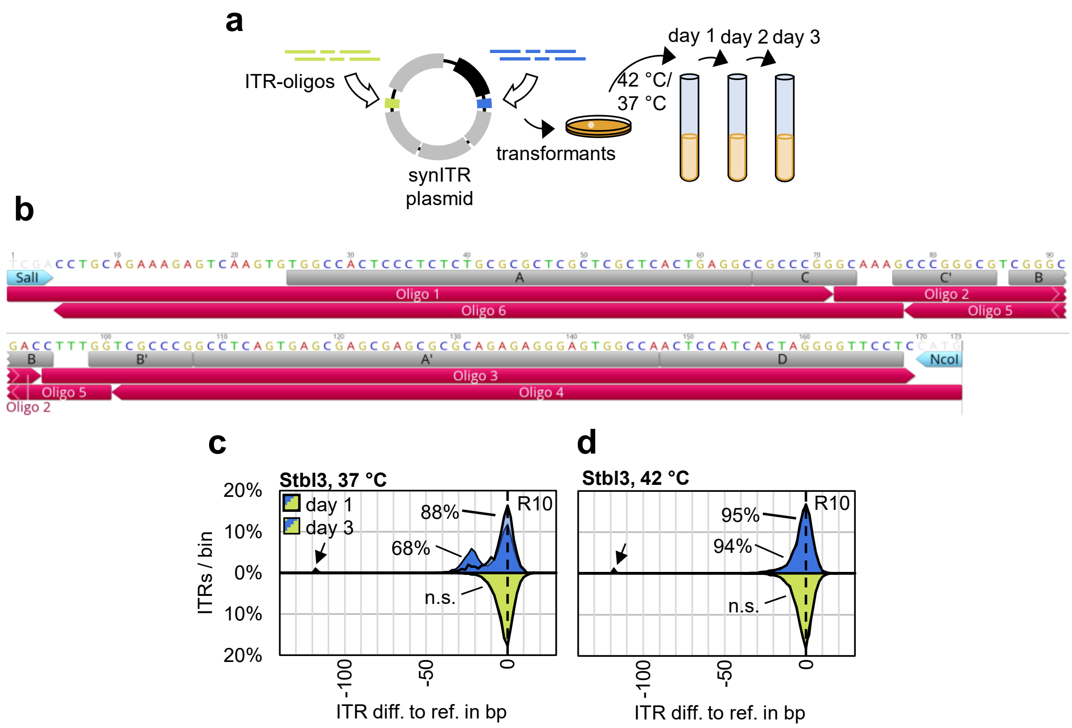

***SI Figure 3:*** *Synthetic ITR cloning (resulting plasmid lab. ref. pZMB990) and degradation in* E. coli*.* ***a****, overview of experimental setup as described in the main method section.* ***b****, details of oligo placement. The oligos (red) were designed such that individual oligos do not form secondary structures as indicated by the annotated ITR hairpin regions (grey). See main methods for cloning details.* ***c****, degradation of the ITRs cloned from oligos in* E. *coli Stbl3 propagated at 37 °C at day one (fat line) and three of propagation as ITR length difference to the reference (144 bp). Percentages indicate percent degradation compared to the upstream ITR at day one. Black arrow: ITRs with complete loss of the palindromic sequence.* ***d****, as (c) but propagation at 42 °C.*

### 6 Sequence alignment of ITR sequences

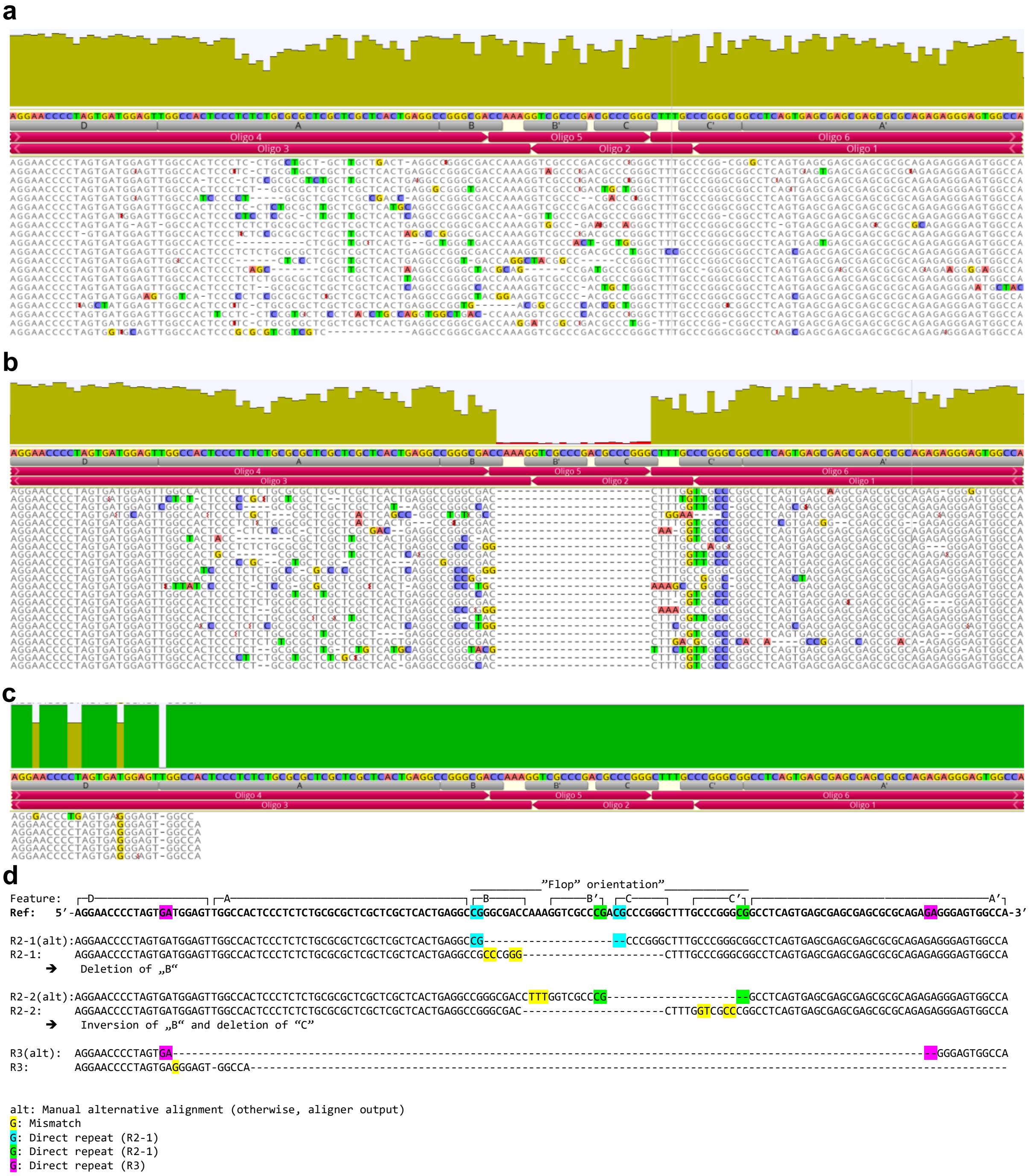

***SI Figure 4:*** *Alignment of ITR noisy raw reads from regions of interest R1 – 3 in* ***Figure 1e*** *(main text) to the AAV2 ITR reference sequence in “FLOP” orientation (top), ori proximal ITR. Sequence features are as originally given in Srivastava, Lusby, and Berns, 1983 (https://doi.org/10.1128/jvi.45.2.555-564.1983). Alignments are given as outputted by the aligner (Geneious) and as manually curated (marked “alt”).* ***a****,* R1 raw ITR sequences comprising full-length ITRs. **b**, R2 raw ITR sequences comprising a mixture of 22 bp deletions with two dominant deletions. **c**, R3 raw ITR sequences comprising mostly the D sequence. **d**, consensus of the dominant sequences of (b) re-aligned to the reference with direct repeats at the junctions highlighted.

### 7 Stabilization of wildtype ITRs in different E. coli strains

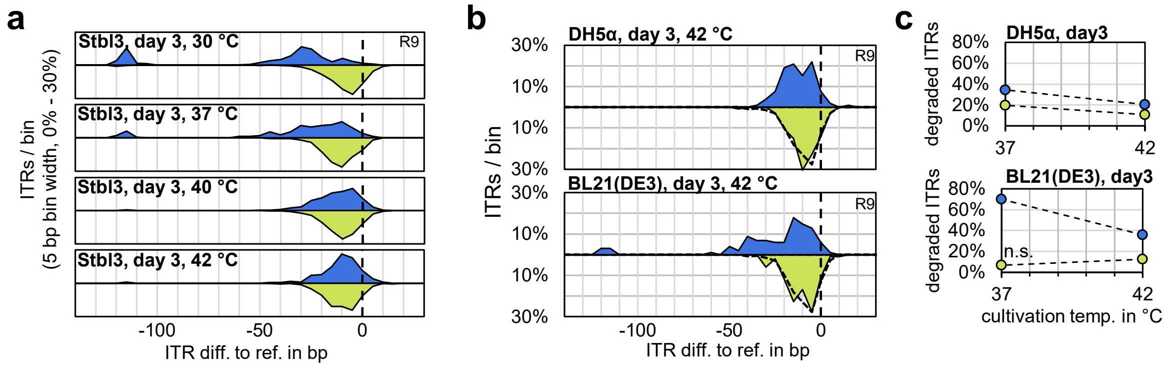

***SI Figure 5:*** *Stabilization of wildtype ITRs (lab. ref. pZMB938) at increased propagation temperature.* ***a****, ITR-length histograms as difference to reference (145 bp) from Stbl3 propagated for three days at the indicated temperatures.* ***b****, similar to (a) but for DH5α and BL21(DE3). Only day three is depicted (dashed line: reference distribution at day one).* ***c,*** *percent degradation over cultivation temperature from the histograms of (a) and (b). R9 data. n.s.: not significant ($p > 0.05$).*

### 8 Investigation of plasmid quality

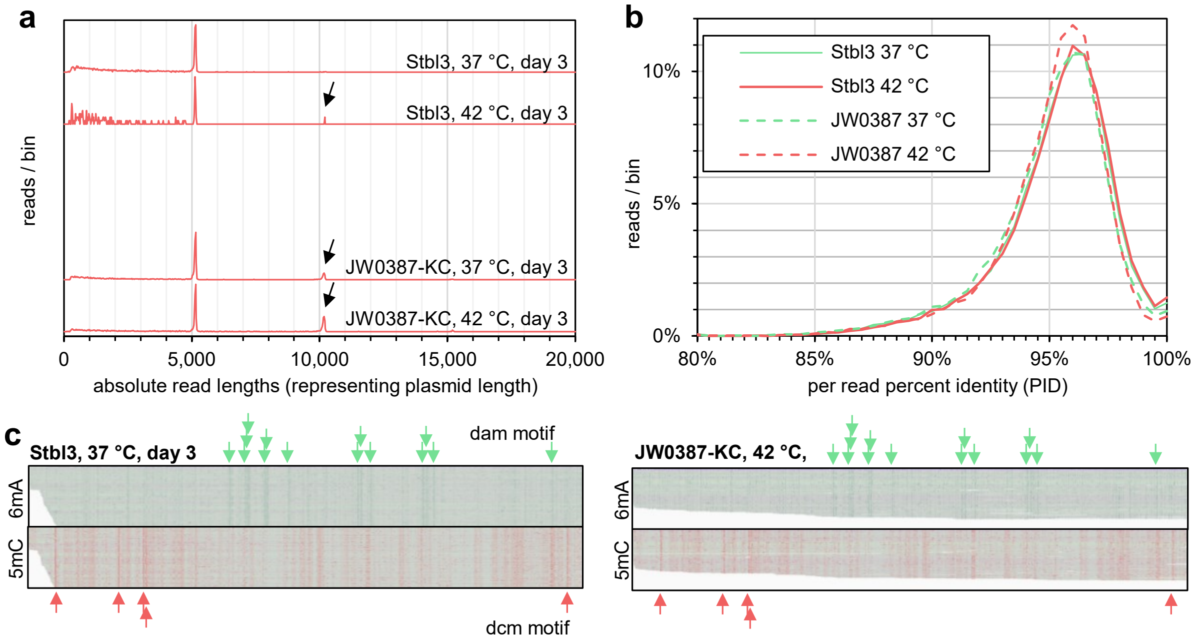

***SI Figure 6:*** *Plasmid quality of an ITR plasmid propagated in Stbl3 and JW0387 at 37 °C and 42 °C.* ***a****, absolute read-length histograms as a measure of plasmid length. A slight increase in concatemerization can be observed at 42 °C propagation compared to 37 °C. Concatemerization is generally more prevalent in JW0387.* ***b****, Histograms of the per-read percent identity of the samples of (A) are visually similar.* ***c****, single-read alignment of reads to the transgene sequence (x-axis) with propensity of modified bases highlighted per position as basis for calculation of 6mA (green) and 5mC methylation (red) of the transgene given in* ***Tabel 1****. Arrows are marking dam and dcm motifs.*

### 9 Slippage model of ITR degradation

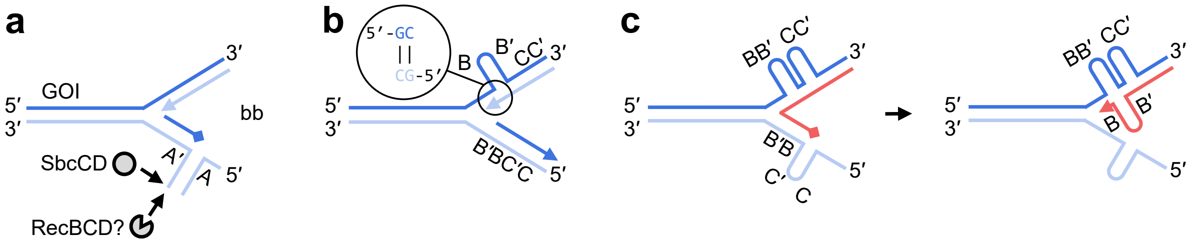

***SI Figure 7:*** *Model for plasmid and ITR sequence loss by slippage.* ***a****, plasmid replication may stall at secondary structures within the replication fork, which are cleaved by SbcCD. Subsequent attack by a processive exonuclease, like RecBCD, may lead to plasmid loss exerting selective pressure on sequences prone to secondary-structure formation.* ***b****, short direct repeats could suffice to bridge an ITR internal hairpin so that plasmid replication can proceed (simple deletion).* ***c****, a more complex inversion-deletion may occur when replication stalls at a secondary structure and switches to the counter-strand, possibly aided by fork regression, before reverting back the original strand for continued replication.*

### 10 Additional proposed pathways to ITR degradation

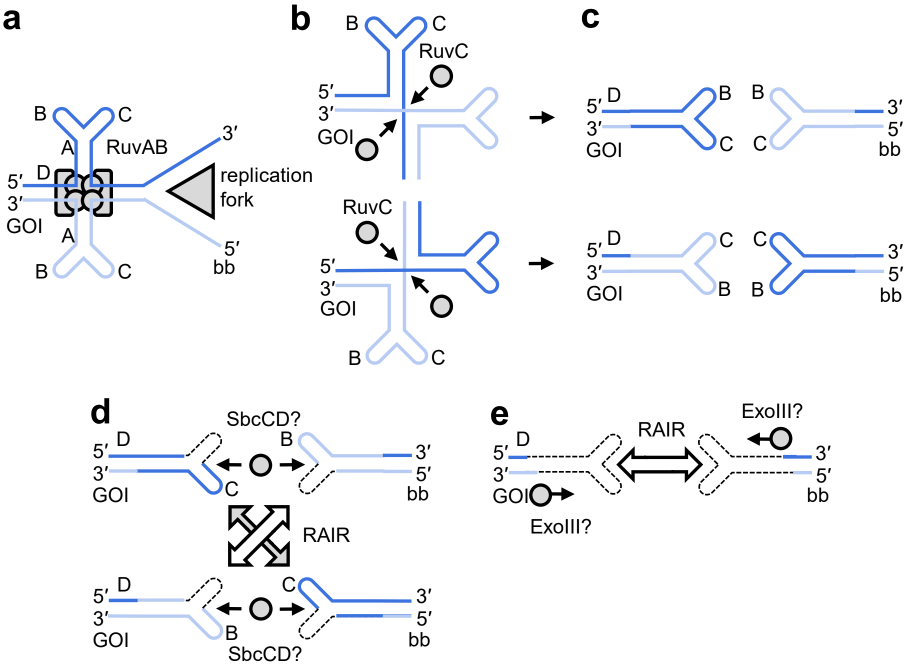

***SI Figure 8:*** *Alternative or aiding model of ITR degradation. The ori proximal ITR is depicted.* ***a****, ITRs may be extruded by negative supercoiling. RuvAB may recognize this structure as a Holliday-like junction and even lock it in place, preventing fork progression.* ***b****, in this case, the structure would be able to isomerize.* ***c****, resolution by RuvC leads to intermediates with back-folded ends containing inverted ITRs.* ***d****, SbcCD or other exonucleases potentially prepare these ends for homologous, RecA-independent repair (RAIR) leading to plasmid rescue.* ***e****, alternatively, exonuclease digest from the point of incision leads to up to a complete palindrome loss and repair by a short homology between the D and A sequence.*

### 11 Bioinformatic analysis of ITR length

We analyzed ITR length in our Nanopore data using a custom Python script. The method takes advantage of the rapid sequencing chemistry, which we have adapted to yield reads that represent the full plasmid, linearized at a random position. The script aligns each read to the plasmid backbone and transgene sequences (without the ITRs), resulting in three alignments per read. The basecalled bases between the backbone and transgene will then be the ITR, of which the length can be calculated and the sequence can be extracted in an unbiased fashion. The principle is illustrated as follows with GOI (transgene, gene of interest without ITRs), bb (backbone), r_st (reference start), r_en (reference end), q_st (query start), q_en (query end):

```
Reference:          .GOI......|.rITR..|..........bb..........|.lITR.|..GOI...
                    ||||||||||        ||||||||||||||||||||||        ||||||||
                   |..........|       |......................|      |........|
 Forward read:     r_st        r_en    r_st                    r_en   r_st     r_en
 Example:          1436        1947     0                      2691    0       1435

 Reverse read:     r_en        r_st    r_en                    r_st   r_en     r_st
 Example:          510          0      2691                     0     1947     511

 Query:            q_st        q_en    q_st                    q_en   q_st     q_en
 Example:           0          511     656                     3618   3758     5195

 Calc. distance:                  145 bp                          140 bp
```

The script utilizes extensive pre-filtering: (1) only reads with three alignments are analyzed further; (2) Only reads with alternating alignments are considered, e.g., GOI-bb-GOI; (3) No ambiguous alignments to the backbone and GOI. For alignments to the forward strand the first (on query) alignment ends within a set number of nucleotides (threshold) of the reference. The second alignment starts and ends within the threshold nucleotides of the reference. The third alignment starts within the threshold nucleotides of the reference. For alignments to the reverse strand the first (on query) alignment starts within the threshold of the reference. The second alignment starts and ends within the threshold of the reference. The third alignment ends within the threshold. The threshold is chosen to be 5 nucleotides. This is illustrated using the following table:

| read | r_st | r_en | strand |
|---|---|---|---|
| 1 | | >max-5 | +1 |
| 1 | <6 | >max-5 | +1 |
| 1 | <6 | | +1 |
| 2 | <6 | | -1 |
| 2 | <6 | >max-5 | -1 |
| 2 | | >max-5 | -1 |

The length of the (two) ITRs per plasmid can be determined individually from the strand-aware sequence of the references as follows (rITR: ori proximal, lITR: ori distal):

| strand | direction | ITRpos |
|---|---|---|
| +1 | GOI->bb | rITR |
| +1 | bb->GOI | lITR |
| -1 | GOI->bb | lITR |
| -1 | bb->GOI | rITR |

The method is functionalized to enable automated analysis of barcoded samples. Initially, the reads are aligned to the two references (backbone and GOI):

```
import mappy as mp
import pandas as pd

def measure_ITRs(ref, sequin):
    a = mp.Aligner(ref, scoring=[2, #Matching Score [2] [default map-ont]
                                 4, #Mismatching penalty [4]
                                 4, #Gap open penalty [4]
                                 2, #Gap extension penalty [2]
                                 24,#Long gap open penalty [24]
                                 1, #Long gap extension penalty [1]
                                 ]) # load or build index
    if not a: raise Exception("ERROR: failed to load/build index")

    data = []
    columns = ["seq_id",   #unique read identifier (given by sequencer)
               "q_en",     #end of alignment on querey (read)
               "seq_len",  #length of read
               "ctg",      #name of reference used for alignment
               "r_st",     #start of alignment on reference
               "r_en",     #end of alignment on reference
               "q_st",     #start of alignment on query
               "strand",   #+1 if on forward stra, -1 if on reverse strand
               "query"]    #original sequence

    for name, seq, qual in mp.fastx_read(sequin): # read a fasta/q sequence
        for hit in a.map(seq):
            tmp = [name, hit.q_en, len(seq), hit.ctg, hit.r_st,
            hit.r_en, hit.q_st, hit.strand, seq]
            data.append(tmp)

    df = pd.DataFrame(data, columns=columns)
    df = df.sort_values(by=["seq_id", "q_st"], ignore_index=True)
    df_grouped = df.groupby(["seq_id", "q_st"], sort=False).last()
```

Then, the DataFrame entries are checked for requirements (1) and (2):

```
#check requirements 1 + 2
    to_drop = []
    for seq_id, hits in df_grouped.groupby(level=0):
        ctg_buff = ()
        if len(hits) == 3: #only hits with three alignments (plasmid monomers)
            for row in hits.iterrows():
                if ctg_buff:
                    if row[1][2] == ctg_buff: #ctg must be alternating
                        to_drop.append(row[0][0])
                        break
                    ctg_buff = row[1][2]
                else:
                    ctg_buff = row[1][2]
        else:
            to_drop.append(seq_id)

    df_grouped.drop(to_drop, axis=0, inplace=True)
```

Adherence to requirement (3) is checked as follows:

```
    #check for requirement 3
    ctg_len = {}
    for name, seq, qual in mp.fastx_read(ref):
        ctg_len[name] = len(seq)

    threshold = 5 #the threshold to determine if an alignment should be disgarded

    to_drop = []
    for seq_id, hits in df_grouped.groupby(level=0):
        idx = 0
        if hits["strand"][0] == 1: #case for forward strand
            for row in hits.iterrows():
                if idx == 0 and row[1][4] < ctg_len[row[1][2]]-threshold:
                    to_drop.append(row[0][0])
                    break
                elif idx == 1 and (row[1][3] > threshold
                or row[1][4] < ctg_len[row[1][2]]-threshold):
                    to_drop.append(row[0][0])
                    break
                elif idx == 2 and row[1][3] > threshold:
                    to_drop.append(row[0][0])
                    break
                idx=idx+1
        else: #case for reverse strand
            for row in hits.iterrows():
                if idx == 0 and row[1][3] > threshold:
                    to_drop.append(row[0][0])
                    break
                elif idx == 1 and (row[1][3] > threshold
                or row[1][4] < ctg_len[row[1][2]]-threshold):
                    to_drop.append(row[0][0])
                    break
                elif idx == 2 and row[1][4] < ctg_len[row[1][2]]-threshold:
                    to_drop.append(row[0][0])
                    break
                idx=idx+1

    df_grouped.drop(to_drop, axis=0, inplace=True)
```

Now, the ITR lengths are determined for the ori proximal ITR (rITR) and ori distal ITR (lITR) and lists containing the individual length are returned for further analysis:

```
    ITR = "rITR"
    ctg_GOI = "GOI"
    ctg_bb = "bb"
    ITR_dict = {"lITR":[1, -1],
                "rITR":[-1, 1]}

    rITR_list = []
    for seq_id, hits in df_grouped.groupby(level=0):
        end_buff = ()

        for row in hits.iterrows():
            if end_buff:
                if ((row[1][2] == ctg_GOI and row[1][5] == ITR_dict[ITR][0]) or
                    (row[1][2] == ctg_bb and row[1][5] == ITR_dict[ITR][1])):
                    rITR_list.append(row[0][1]-end_buff)
                end_buff = row[1][0]
            else:
                end_buff = row[1][0]

    ITR = "lITR"
    ctg_GOI = "GOI"
    ctg_bb = "bb"
    ITR_dict = {"lITR":[1, -1],
                "rITR":[-1, 1]}

    lITR_list=[]
    for seq_id, hits in df_grouped.groupby(level=0):
        end_buff = ()

        for row in hits.iterrows():
            if end_buff:
                if ((row[1][2] == ctg_GOI and row[1][5] == ITR_dict[ITR][0]) or
                    (row[1][2] == ctg_bb and row[1][5] == ITR_dict[ITR][1])):
                    lITR_list.append(row[0][1]-end_buff)
                end_buff = row[1][0]
            else:
                end_buff = row[1][0]

    return (rITR_list, lITR_list)
```

Alternatively, should the respective ITR sequence instead of the length be of interest, it is possible to modify the last for-loop so that “ITR_list” contains the sequences:

```
for seq_id, hits in df_grouped.groupby(level=0):
end_buff = ()

for row in hits.iterrows():
    if end_buff:
        if ((row[1][2] == ctg_GOI and row[1][5] == ITR_dict[ITR][0]) or
            (row[1][2] == ctg_bb and row[1][5] == ITR_dict[ITR][1])):
            rITR_list.append((row[0][1]-end_buff,
                                     row[1][6][end_buff:row[0][1]]))
            labels.append(len(row[1][6][end_buff:row[0][1]]))
        end_buff = row[1][0]
    else:
        end_buff = row[1][0]
```

### 12 Plasmid maps

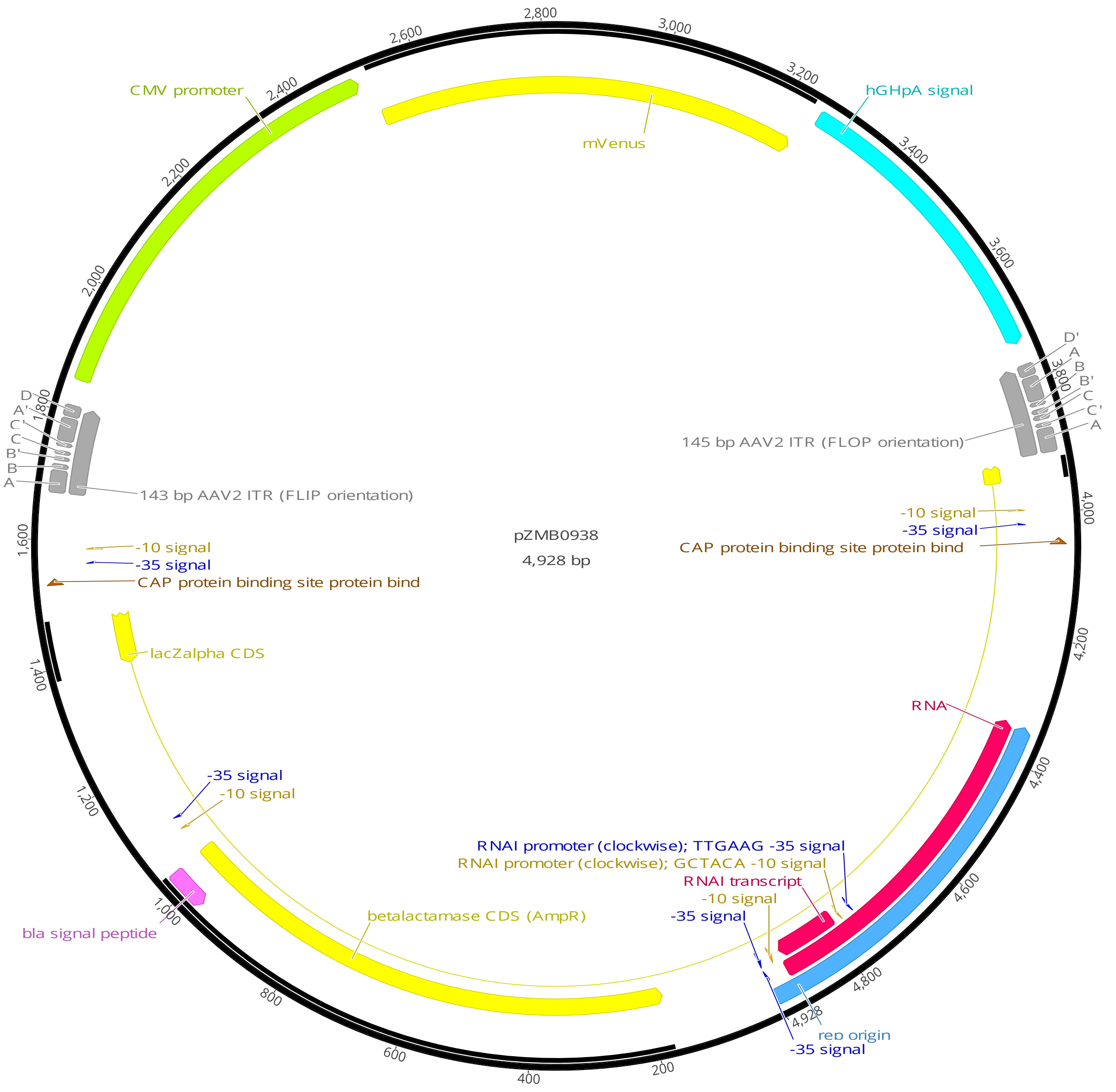

***SI Figure 9:*** *Plasmid map of pZMB938, a pUC19-based plasmid carrying an mVenus fluorescence reporter flanked by a 143 bp ITR (FLIP orientation) and a 145 bp ITR (FLOP orientation). For plasmid construction, see main methods.*

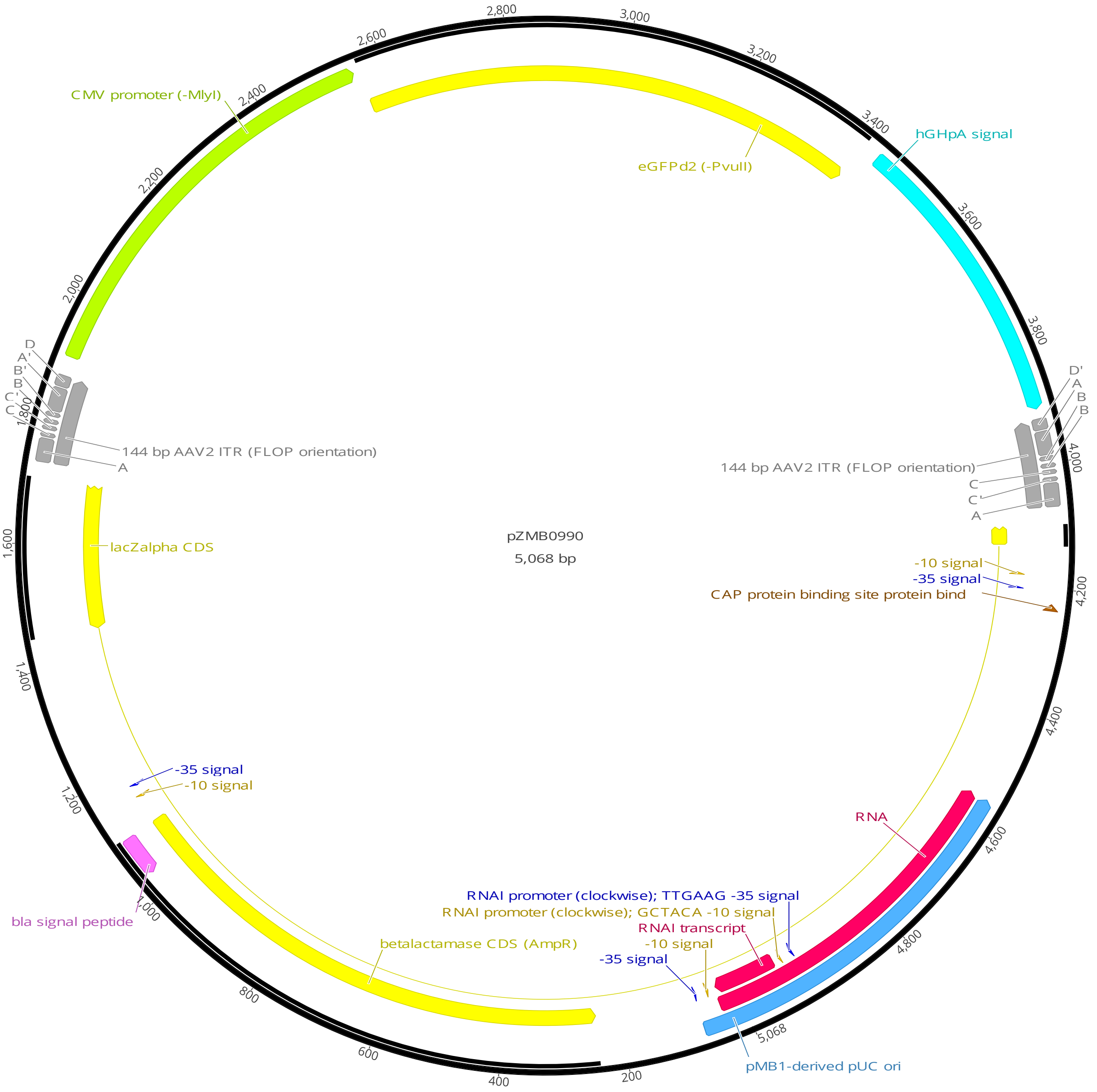

***SI Figure 10:*** *Plasmid map of pZMB990, a pUC19-based plasmid carrying an eGFPd2 fluorescence reporter flanked by two 144 bp ITRs (FLOP orientation). For plasmid construction, see main methods.*
